# Establishment of murine hybridoma cells producing antibodies against spike protein of SARS-CoV-2

**DOI:** 10.1101/2020.08.29.272963

**Authors:** Nadezhda V. Antipova, Tatyana D. Larionova, Michail I. Shakhparonov, Marat S. Pavlyukov

## Abstract

In 2020 the world faced the pandemic of COVID-19 - severe acute respiratory syndrome caused by a new type of coronavirus named SARS-CoV-2. To stop the spread of the disease, it is crucial to create molecular tools allowing to investigate, diagnose and treat COVID-19. One of such tools are monoclonal antibodies (mAbs). In this study we describe the development of hybridoma cells that can produce mouse mAbs against receptor binding domain of SARS-CoV-2 spike (S) protein. These mAbs are able to specifically detect native and denaturized S protein in all tested applications including immunoblotting, immunofluorescence staining and enzyme-linked immunosorbent assay. In addition, we showed that the obtained mAbs decreased infection rate of human cells by SARS-CoV-2 pseudovirus particles in *in vitro* experiments. Finally, we determined the amino acid sequence of light and heavy chains of the mAbs. This information will allow to use the corresponding peptides to establish genetically engineered therapeutic antibodies. To date multiple mAbs against SARS-CoV-2 proteins have been established, however due to the restrictions caused by pandemic, it is imperative to have a local source of the antibodies suitable for researches and diagnostics of COVID-19. Moreover, as each mAb has a unique binding sequence, bigger sets of various antibodies will allow to detect SARS-CoV-2 proteins even if the virus acquires novel mutations.

## Introduction

At the beginning of 2020 the world faced an outbreak of COVID-19 - severe acute respiratory syndrome caused by SARS-CoV-2 coronavirus. [1, 2]. More than 20 million people were infected during the first 8 months of the pandemic. To slow down the spread of the disease WHO put significant effort into supporting scientific research and the development of diagnostics, vaccines and medications against COVID-19 [3], multiple platforms were established to monitor real time distribution of the disease all over the world [4].

Phylogenetic analysis has attributed SARS-CoV-2 to the genus of Betacoronavirus in the Coronaviridae family [5, 6]. Despite the name and the genetical similarity, SARS-CoV-2 is not a direct descendant of previously described SARS-CoV virus. Rather it has independently originated during the evolution [7]. Growing number of full genome sequences of SARS-CoV-2 have revealed multiple mutations and deletions in coding and non-coding regions of the virus [8]. However, SARS-CoV-2 has a relatively low mutation rate due to high accuracy of the enzymes involved in virus replication [9].

The first step of coronavirus entry into the cell is its interaction with the surface receptors of the host. These receptors include angiotensin-converting enzyme 2 (ACE2) for SARS-CoV and SARS-CoV-2 [10–12] and CD26 for MERS-CoV [13]. Spike (S) glycoprotein is responsible for the interaction of SARS-CoV-2 with ACE2. This protein contains a receptor binding domain (RBD) that interacts with the N-terminal peptidase domain of ACE2 with Kd of 14.7 nM [14–16]. Cryogenic electron microscopy has revealed a molecular structure of ACE2*RBD protein complex [14, 15]. Due to its’ important role and a unique structure, RBD is considered as one of the main targets for the development of neutralizing antibodies against SARS-CoV-2.

In addition to the mentioned above, S protein of SARS-CoV-2 contains peptide sequences that may bind to MHC and serve as an effective epitope for the generation of antibodies [17]. It was previously shown that S and N proteins of SARS-CoV induce prominent and prolonged T-cell immune responses [18], which is consistent with recent observation for SARS-CoV-2 [19]. With its currently evolving role S protein might serve as a promising antigen for the development of vaccines against SARS-CoV-2 [20]. To this date, more than 20 vaccines based on the S protein are undergoing clinical trials [21].

Besides the vaccines, development of diagnostics is critically important to stop the spreading of COVID-19. Majority of tests for SARS-CoV-2 are being performed using reverse transcription polymerase chain reaction (PCR) on swabs from the nasopharynx or upper respiratory tract. Additionally, computed tomography of the chest is used to confirm the diagnosis, but its results are nonspecific and can often coincide with other diseases, therefore the diagnostic value of this method is limited [22, 23]. Serological tests of the immune response of patients are also important as presence of specific antibodies allows to determine the prevalence of COVID-19 in the society and identify people who can potentially be immune to the infection. It was also proposed to use an immunoassay to evaluate the response to the vaccination [24, 25]. Although the presence of neutralizing antibodies can only be confirmed by specific tests using virus-like particles [26], high titers of IgG detected by immunoassay have been shown to positively correlate with the amount of neutralizing antibodies. To avoid the emergence of virus resistance to the antibodies, administration of cocktails containing multiple therapeutic antibodies was proposed as a treatment strategy [27, 28].

Currently the manufacturing of antibodies shares one of the leading places on the marked together with the production of vaccines [29]. The number of mAbs based drugs approved for the clinic is growing every year. In addition to human or humanized neutralizing antibodies proposed as a “magic bullet” for the treatment of COVID-19 [30], there is now a high variety of antibodies produced in different species against all proteins of the SARS-CoV-2. These antibodies cannot be directly used to treat patients with COVID-19, but they serve as an important tool for both basic scientific research and for the development of vaccines and diagnostic kits for SARS-CoV-2. Unfortunately, none of these antibodies were developed in the Russian Federation. However, during the pandemic it is crucial to have a local source of the antibodies suitable for researches and diagnostics of COVID-19.

In the present study we describe the development of hybridoma cells that produces mAbs against RBD of SARS-CoV-2 S protein. We also performed extensive characterization of the obtained mAbs and describe their usage for various applications.

## Experimental procedures

### Plasmid Construction

The DNA fragment encoding RBD of SARS-CoV2 was amplified by PCR using primers S2_for (AAA AGC TAG CAA TGG CAC GAT AAC TGA CGC) and S2_rev (AAT TAA GCT TAA ACA CAT TTG ACC CAG TTG AGT A) from pTwist-EF1a-nCoV-2019-S-2xStrep plasmid kindly provided by Dr. Nevan J. Krogan [31]. Resulting DNA fragment was cloned into NheI/HindIII sites of pET28a+ plasmid (Novagen) to generate pET28-S plasmid. The absence of unwanted mutations in the inserts and vector-insert boundaries was verified by sequencing.

### Recombinant protein expression and purification

To produce RBD fragment of S protein in bacterial cells BL21 (DE3) Codone+ RIL E. coli cells (Agilent) were transformed with pET28-S plasmid. Bacteria were incubated at 37°C on shaker until OD_600_ reached 0.7. Next, IPTG was added to a final concentration 1mM and bacteria were incubated for an additional 4 hours at 37°C. 200 ml media with bacteria were centrifuged for 15 min, 5000 g at 4°C and the pellet was resuspended in 12 ml of lysis buffer B (200 mM NaCl, 100 mM NaH_2_PO_4_, 10 mM Tris-HCl pH 8.0, 8 M Urea, 0.5 mM DTT) and incubated for 1,5 hours at room temperature. Next, solution was centrifuged for 15 min, 20000 g at 4°C and supernatant was incubated with 2 ml of Ni-NTA resin for 1 hour under a constant agitation. Suspension was transferred to a column and washed with 20 ml of buffer B and 10 ml of buffer C (same as buffer B but pH 6.3). Bounded proteins were eluted with buffer D (buffer C with 250 mM of imidazole) and dialyzed overnight against PBS with 1mM DTT. The purity of obtained protein was assessed by electrophoresis and subsequent Coomassie Blue staining. RBD fragment of S protein purified from HEK293 cells was kindly provided by Vassili N. Lazarev from the Center for Precision Genome Editing and Genetic Technologies for Biomedicine, Federal Research and Clinical Center of Physical-Chemical Medicine of Federal Medical Biological Agency.

### Cell Culture

HEK293 and HT1080 cells were grown in Dulbecco’s modified Eagle’s medium (DMEM) supplemented with 10% (v/v) fetal bovine serum (FBS), 2mM L-glutamine, 1mM Na-pyruvate and penicillin-streptomycin mixture (100μg/ml). Transfection was performed with Lipofectamine LTX reagent (Thermo Fisher) according to the manufacturer’s protocol. Transfected cells were stained or injected into mice 48 hours after transfection. X63 myeloma and hybridoma cells were grown in DMEM/F12 medium supplemented with 15% (v/v) FBS, GlutaMAX (Thermo Fisher), 1mM Na-pyruvate and penicillin-streptomycin mixture (100μg/ml). HAT or HT supplements (Sigma) were added at different time points after fusion as described previously [32].

### Mice immunization and hybridoma fusion

5 weeks old BALB/c mice were immunized according to the standard protocol [32] with several modifications. First immunization was performed intraperitoneally with 200μg of RBD resuspended in 150μl of PBS and mixed with 150μl of either Freund’s complete adjuvant (FCA) or Freund’s incomplete adjuvant (FIA) (Thermo Fisher). Two weeks later mice were intraperitoneally injected with 200μg of RBD resuspended in 150μl of PBS and mixed with 150μl FIA. 6 days later blood from the tail vein was collected to perform ELISA and IF staining. At days 28, 29 and 30 after first immunization mice were subcutaneously injected with 2*10^6^ HEK293 cells transfected with pTwist-EF1a-nCoV-2019-S-2xStrep plasmid. At day 40 mice were intraperitoneally injected with 100μg of RBD resuspended in 100μl of PBS and 4 days later mice were sacrificed and splenocytes were isolated and subsequently fused with X63 myeloma cells according to the standard protocol [32]. All animal experiments were carried out under an Institutional Animal Care and Use Committee (IACUC) approved protocol according to NIH guidelines.

### Production of ascites

Ascites were prepared to obtain large quantities of mAbs [32]. Briefly, mice were intraperitoneally injected with 200μl of FIA. Five days later, mice were intraperitoneally injected with 2.5*10^6^ of hybridoma cells in 150μl of PBS. After 12 days, mice were sacrificed and ascitic fluid was harvested from the intraperitoneal cavity and centrifuged at 20 000g at 4 °C for 15 min.

### Enzyme-linked immunosorbent assay

Enzyme-linked immunosorbent assay (ELISA) was performed as described previously [33]. Briefly, 0.5 μg of RBD isolated from E. coli; RBD isolated from or HEK293 cells or 0.5 μg of a control protein isolated from E. coli were immobilized on wells of 96 well EIA plate (Corning). After blocking with 1% BSA in TBST (20 mM Tris pH 7.6, 150 mM NaCl, 0.1% Tween20) wells were incubated with culture medium from hybridoma cells diluted 1:1 in TBST; mouse serum diluted 1:700 in TBST; or ascitic fluid diluted in TBST. After washing with TBST wells were incubated with HRP-conjugated anti-mouse secondary antibodies (Thermo Fisher) (1:10000 dilution in TBST) and developed with 1-Step Ultra TMB-ELISA Substrate Solution (Thermo Fisher) according to the manufacturer’s protocol.

### Immunoblotting

Immunoblotting (WB) was performed as described previously [34]. Briefly, cells were lysed in RIPA buffer (150 mM NaCl, 1% NP40, 0.5% Na-deoxycholate, 0.1% SDS, 50 mM Tris pH 8.0, Protease inhibitor cocktail (Sigma)) on ice for 30 min and then centrifuged at 18 000g, 4°C for 10 min. Supernatant was used for SDS-PAGE. Alternatively, cells were lysed by boiling at 100°C for 10 minutes in 0.5% SDS and subsequently used for deglycosylation and SDS-PAGE. After electrophoreses proteins were transferred to PVDF membrane and incubated overnight with culture medium from hybridoma cells diluted 1:1 in TBST. After washing membrane was incubated with HRP-conjugated anti-mouse secondary antibodies (Thermo Fisher) (1:4000 dilution in TBST with 5% nonfat dry milk). Membranes were developed with SuperSignal West Pico PLUS chemiluminescent substrate (Thermo Fisher) and analyzed on ImageQuant LAS 500 imager (GE Healthcare).

### Deglycosylation

Deglycosylation of S protein from the lysate of HEK293 cells transfected with pTwist-EF1a-nCoV-2019-S-2xStrep plasmid was performed as described previously [35] using PNGase F (New England Biolabs).

### Immunofluorescence microscopy

HT1080 cells were plated in wells of Lab-Tek II chamber and co-transfected with pTwist-EF1a-nCoV-2019-S-2xStrep and pTagGFP2-C (Evrogen) plasmids. Two days after transfection cells were washed with phosphate buffered saline (PBS) and fixed with 4% PFA in PBS for 15 min at room temperature. Cells were washed 2 times with PBS and incubated with culture medium from hybridoma cells diluted 1:1 in PBS or with mouse serum diluted 1:700 in PBS. After 5 washes with PBS cells were incubated with AlexaFluor555-conjugated anti-mouse secondary antibodies (Thermo Fisher) (1:500 dilution in PBS) and subsequently stained with DAPI. Images were captured with a DIAPHOT 300 fluorescent microscope (Nikon).

### Generation of SARS-CoV2 pseudovirus particles

HEK293FT cells in T75 flask were cotransfected with 6 μg of pTwist-EF1a-nCoV-2019-S-2xStrep plasmid, 9 μg of psPAX2 plasmid and 12 μg of pCDH-GFP-IRES-Puro plasmid [36]. Next day culture media was changes and 72 hours later media with viruses was collected and filtered through 0.4 μm filter. Pseudoviruses were concentrated by ultracentrifugation as described previously [37].

### Analysis of infection efficiency

HT1080 cells were plated in wells of 24 well plate to 50% confluence. 180 μl of fresh media, 180 μl of hybridoma media (with or without Abs) and 40 μl of virus suspension were added to each well. 4 days later cells were extensively washed with PBS and genomic DNA was isolated using Lira reagent (Biolabmix) according to the manufacturer’s protocol. Obtained DNA was used for real-time PCR with primers for 18S (GGC CCT GTA ATT GGA ATG AGT C and CCA AGA TCC AAC TAC GAG CTT) and GFP (TGG TGA CCA CCC TCT GCT AC and GGC GCT CTT GAA GAA GTC GT). PCR was performed on LightCycler 96 Instrument (Roche) as described previously [36].

### Antibody sequencing

Sequencing of mRNAs encoding heavy and light chains of antibodies was performed using 5’ SMART RACE method as described previously [38] with slight modifications. First strand of cDNA was synthesized using Mint kit (Evrogen) with primers for k-chain (TTG TCG TTC ACT GCC ATC AAT C), λ-chain (GGG GTA CCA TCT ACC TTC CAG), and heavy-chain (CTG GAC AGG GAT CCA GAG TTC CA) according to the manufacturer’s protocol. Next, cDNA encoding immunoglobulins was amplified by conventional PCR using M1 forward primer (AAG CAG TGG TAT CAA CGC AGA GT) and revers primers specific for k-chain (ACA TTG ATG TCT TTG GGG TAG AAG), λ-chain (ATC GTA CAC ACC AGT GTG GC), and heavy-chain (GGG ATC CAG AGT TCC AGG TC). Obtained DNA was purified and cloned into pKAN-T (Evrogen) vector that was subsequently sequenced. At least 3 different clones were sequenced for each chain of the antibodies. Immunoglobulin sequences were analyzed using IgBLAST (https://www.ncbi.nlm.nih.gov/igblast/), BLASTn (https://blast.ncbi.nlm.nih.gov/Blast.cgi?PROGRAM=blastn&PAGE_TYPE) and SignalP-5.0 (http://www.cbs.dtu.dk/services/SignalP/) software.

## Results

### Mice immunization with S protein of SARS-CoV2

Animal immunization is the first step in the development of mouse mAbs. There are multiple proposed protocols to create antibodies against different types of proteins and each of them has certain benefits and drawbacks. Thus, immunization with DNA vectors or cells overexpressing target protein on their surface allows production of antibodies against native protein which has correct folding and proper post-translational modifications [39, 40]. On the other hand, immunization with synthetic peptides or recombinant protein fragments enables the development of antibodies specific for a certain part of amino acid sequence without contamination of antibodies raised against oligosaccharides decorating the protein [41]. In the current study, we applied combined protocol and first immunized animals with recombinant RBD fragment purified from E. coli and next injected mice with cells overexpressing full length S protein.

On the first step, we expressed and isolated RBD of S protein from E. coli under denaturing conditions (**Fig. 1A**). Despite multiple attempts, we were unable to design a refolding protocol that could allow us to obtain a soluble protein in a buffer suitable for immunization. Therefore, we injected mice with a suspension of insoluble protein in PBS. The schematic representation of immunization schedule is shown on **figure 1B**. First immunization was performed with recombinant protein mixed at a 1:1 ratio with either Freund’s complete adjuvant (FCA) or Freund’s incomplete adjuvant (FIA). Interestingly, FIA allowed to obtain a significantly higher titer of specific antibodies as opposed to FCA (**Fig. 1C**). This result indicates that insoluble RBD is highly immunogenic and does not require any stimulating additives for the development of immune response. Similar data were obtained previously during animal immunization with the surface protein of the Zika virus [41].

**Fig. 1.**
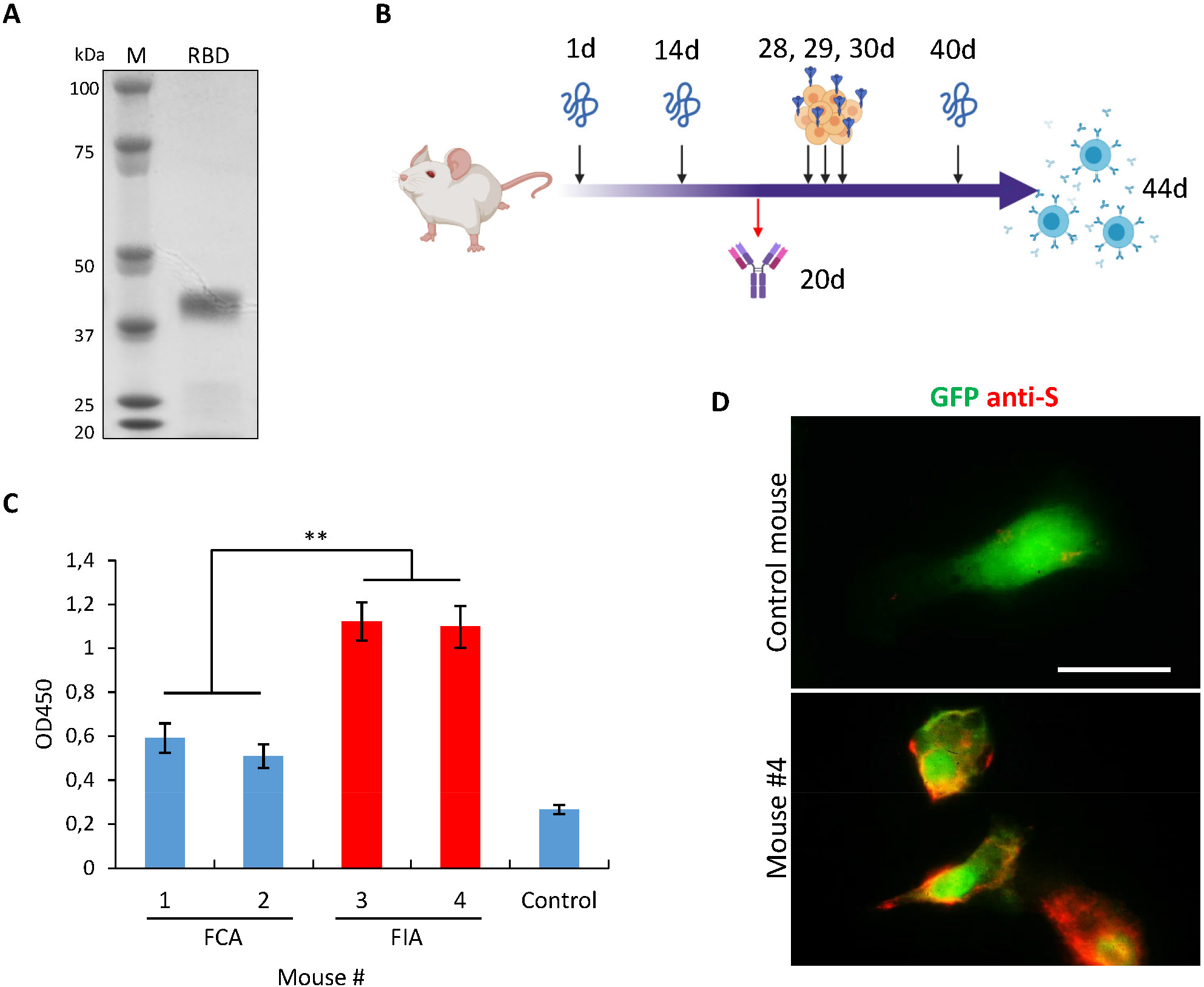
Immune responses of mice injected with SARS-CoV-2 S protein. *A*, Electrophoresis of recombinant RBD that was purified from E. coli. *B*, Schematic representation of mice immunization workflow. At days 1, 14 and 40 mice were injected with RBD purified from E. coli; at day 20 blood was collected to test the presence of Abs against RBD; at days 28, 29 and 30 mice were injected with HEK293 cells overexpressing full length S protein; at day 44 mice were sacrificed to isolate splenocytes. *C*, ELISA showing humoral immune responses of mice immunized with RBD suspension mixed 1:1 with FCA or FIA; non-immunized mouse was used as a control. *D*, Fluorescence images of cells cotransfected with plasmids encoding GFP (green) and full length S protein and subsequently stained with serum obtained from mice #4 (red). **p < 0.01.

Next, Immunofluorescence staining (IF) of cells overexpressing full length S protein with mice serum demonstrated that antibodies raised against insoluble RBD could detect native S protein (**Fig. 1D**). Therefore, both insoluble RBD and full length S protein expressed on the surface of mammalian cells have common epitopes indicating that these proteins could be used for the production and selection of antibodies against SARS-CoV2.

### Selection of hybridoma cells secreting mAbs against S protein of SARS-CoV2

44 days after the first immunization splenocytes were purified from the mice and subsequently fused with X63 myeloma cells. After the fusion cells were seeded on 20 96-wells plates and 3 weeks later 120 hybridoma monoclones were obtained. mAbs produced by these clones were tested by enzyme-linked immunosorbent assay (ELISA) against RBD purified from HEK293 cells (**Fig. 2A**). Primary screening with the protein that was purified from the different sources (compared to the protein used for immunization) allowed us to exclude clones that produced antibodies against various contaminants that inevitably present in the samples used for immunization. It is important to note that RBD expressed in HEK293 cells presumably has glycosylation pattern which is similar to that of native S protein and therefore mAbs interacting with RBD from HEK293 most likely could interact with full length S.

**Fig. 2.**
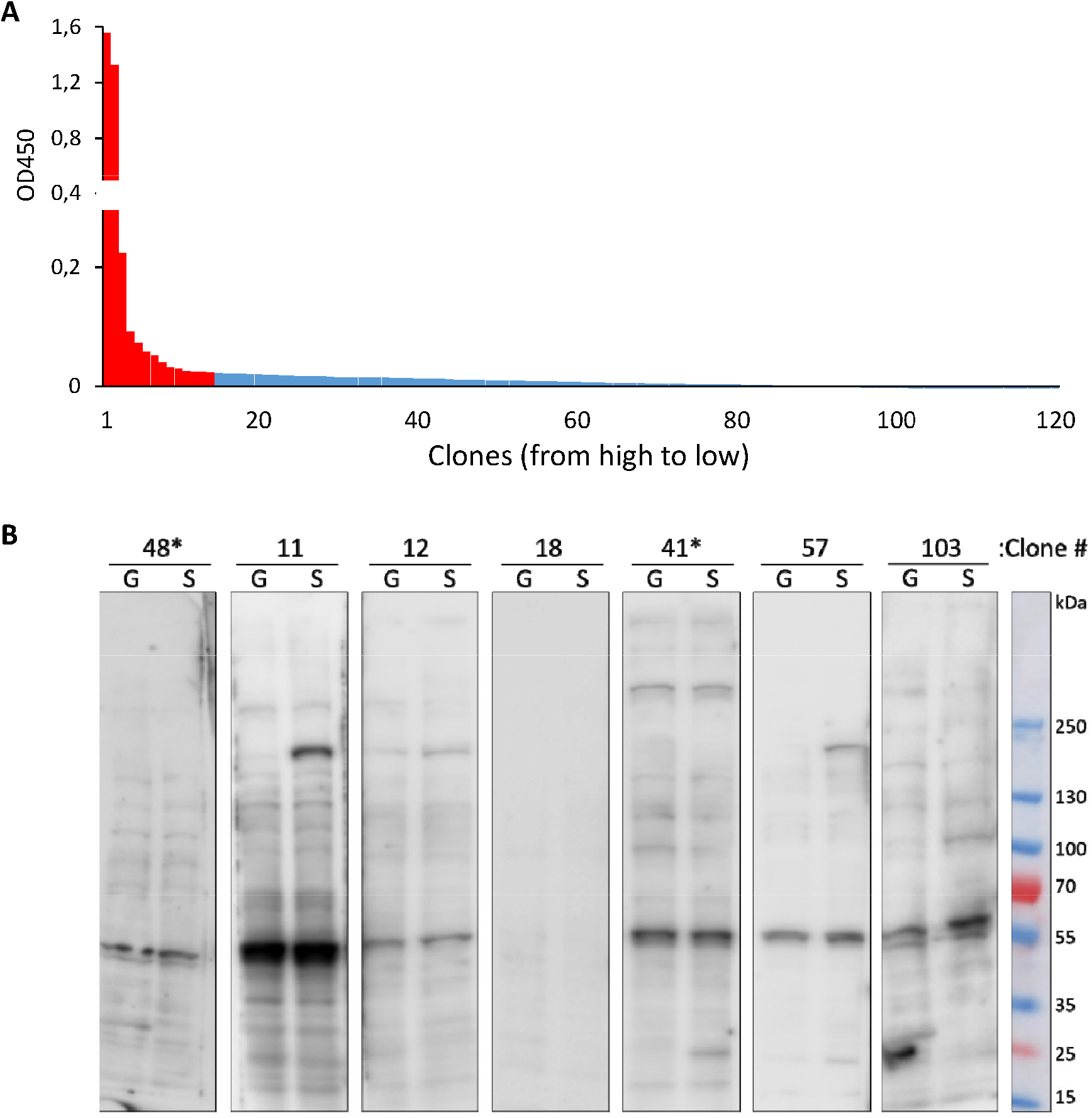
Characterization of Abs secreted by hybridoma monoclones that were obtained after fusion. *A*, ELISA showing reactivity of Abs from 120 different monoclones to RBD purified from HEK293 cells; red-indicates clones that were used for further analysis. *B*, Representative immunoblots of cells transfected with plasmid encoding GFP (G) or full length S protein (S). Culture medium from different monoclones were used to stain the membranes. * indicates clones with highest reactivity according to ELISA results.

Total of fourteen hybridomas that demonstrated the strongest immunoreactivity against RBD were chosen for further analysis. We performed immunoblotting (WB) with the culture medium from our clones to detect S protein in the lysates of human cells that were transfected with plasmids encoding full length S protein or GFP as a control. Representative WBs are shown on **figure 2B**. Interestingly, two clones which had highest immunoreactivity in ELISA experiment failed to detect S protein on WB. Therefore, mAbs produced by these cells are probably nonspecific. Another important observation is that most of the tested mAbs were able to detect endogenous human protein with molecular weight of approximately 60 kDa. It is possible that this protein may have a common epitope with RBD of S protein from SARS-CoV2.

### Establishment of monoclones producing antibodies against S protein of SARS-CoV2

It is well known that hybridoma cells are characterized by the high degree of genetical instability during the first passages. Thus, descendants of a single hybridoma cell will most likely produce different antibodies at early time points after fusion [42]. Therefore, we chose hybridoma #11 which showed highest immunoreactivity against full length S protein (**Fig. 2B**) and subcloned these cells to obtain true monoclones. Culture media from 17 subclones of hybridoma #11 were tested by ELISA against RBD purified from E. coli, RBD purified from HEK293, and control protein purified from E. coli (**Fig. 3A**). Based on our results, we picked 3 subclones for further analysis. Subsequent WB analysis demonstrated that all monoclones produce mAbs with higher specificity compared to the original hybridoma #11 (**Fig. 3B**). It is interesting to note that mAbs from monoclone #11/13 which showed highest signal against RBD purified from E. coli but not against RBD purified from HEK293 (**Fig. 3A**) were only able to detect endogenous 60 kDa human protein and showed weak binding to full length S protein. Therefore, it is possible that recombinant insoluble RBD has high epitope similarity to the undetermined endogenous protein.

**Fig. 3.**
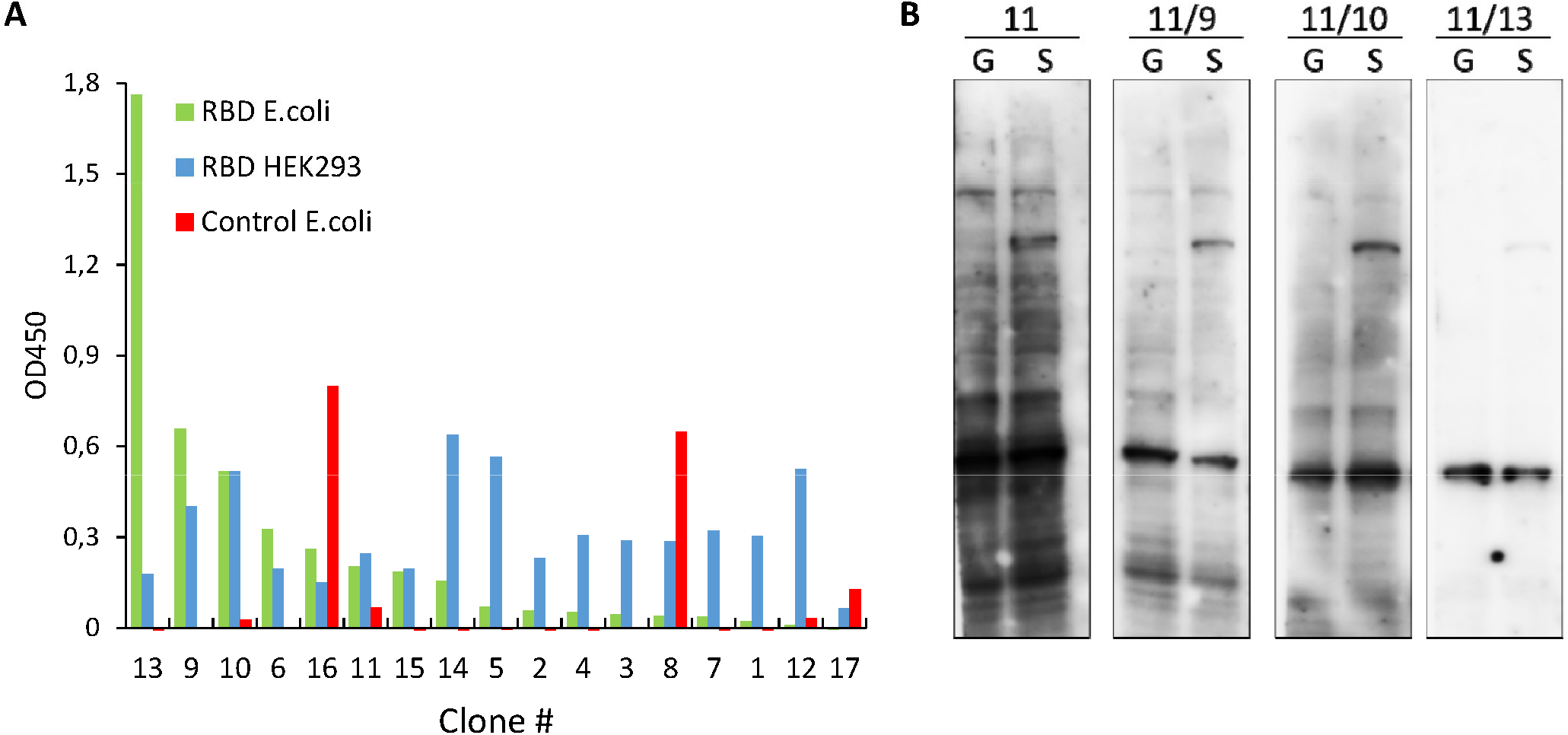
Characterization of mAbs secreted by monoclones that were obtained after subcloning of hybridoma #11. *A*, ELISA showing reactivity of mAbs from 17 different subclones to RBD purified from E. coli (green), RBD purified from HEK293 cells (blue) and to control protein purified from E. coli (red). *B*, Immunoblots of cells transfected with plasmid encoding GFP (G) or full length S protein (S). Culture medium from different subclones were used to stain the membranes.

### Characterization of mAbs against S protein of SARS-CoV2

Based on our results (**Fig. 3**) we chose mAbs produced by monoclone #11/9 for further characterization. First, we used immunofluorescence staining to test if these mAbs were able to bind to native nondenatured S protein. Images on **figure 4A** shows that mAbs #11/9 specifically stain human cells that overexpress S protein and have little or no binding to the neighboring nontransfected cells. Therefore, mAbs #11/9 were able to detect RBD fragment in ELISA experiment; full length denatured S protein in WB; and full length native S protein on a cell surface as demonstrated by IF microscopy.

**Fig. 4.**
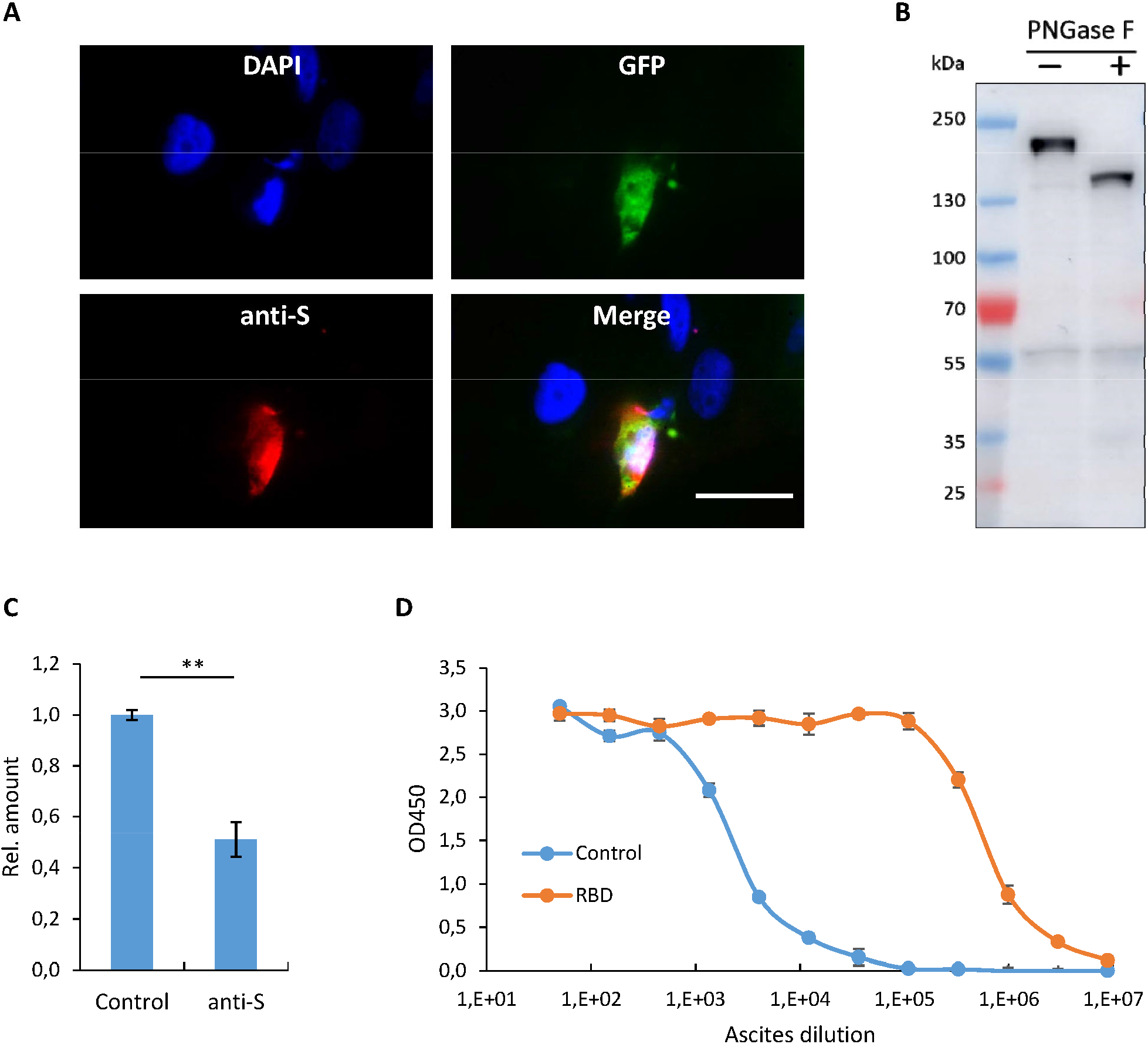
Characterization of mAbs secreted by monoclone 11/9. *A*, Fluorescence images of cells cotransfected with plasmids encoding GFP (green) and full length S protein and subsequently stained with DAPI (blue) and mAbs secreted by monoclone #11/9 (red). *B*, Immunoblot analysis of cells transfected with plasmid encoding full length S protein and lysed by boiling in SDS. Lysate was incubated in a presence or absence of PNGase F. Membrane was stained with mAbs #11/9. *C*, Relative incorporation level of GFP gene into the genome of HT1080 cells infected with SARS-CoV-2 pseudovirus particles encoding GFP. Infection was performed in presence or absence of culture medium from #11/9 hybridoma cells. *D*, Serial dilution ELISA showing reactivity of ascites formed by monoclone #11/9 against RBD (red) or control protein (blue) purified from E. coli. **p < 0.01.

It was previously shown that RBD undergoes extensive glycosylation in human cells [43]. Therefore, we aimed to test if glycosylation affects binding of mAbs #11/9 to the S protein. To this end, we treated lysate of cells overexpressing S protein with deglycosylating enzyme PNGase F. **Figure 4B** demonstrates that deglycosylation decreases molecular weight of S protein by at least 30 kDa, however it has no effect on the intensity of staining with mAbs #11/9. Thus, it is reasonable to conclude that the region of S protein that interacts with mAbs #11/9 is not masked by glycoside groups of the protein.

RBD plays the key role in the virus infection by interacting with the surface protein ACE2. Hence antibodies against RBD can effectively block infection of human cells with SARS-CoV2 [26]. To test the effect of mAbs #11/9 on the infection rate, we obtained SARS-CoV2 pseudovirus particles that encode GFP. We used these particles to transduce HT1080 cells which were previously shown to be highly susceptible for the infection with SARS-CoV [44]. Unfortunately, we were unable to reach more than 5% transduction efficiency. Therefore, to quantify the effect of mAbs #11/9 on the infection rate, we determined the level of DNA encoding GFP that was incorporated into the genome of HT1080 cells after the transduction with pseudoviruses. Our results demonstrate that the presence of mAbs #11/9 decreases the infection rate of SARS-CoV2 pseudovirus particles by at least two folds (**Fig. 4C**).

Finally, we aimed to produce mAbs #11/9 in higher quantities and test how these mAbs will recognize S protein when used in the different concentrations. For this reason, we injected two mice with corresponding hybridoma cells and 12 days later both animals formed ascites. In total 6 ml of ascitic fluid was collected. We used this ascites for serial dilution ELISA with RBD purified from E. coli. According to our data, mAbs #11/9 from ascites were able to detect recombinant RBD but not a control protein in as high as 1:1000 000 dilution (**Fig. 4D**), indicating an efficient binding of the antibodies to the antigen.

### Identification of amino acid sequences of mAbs #11/9

During last few decades genetically modified antibodies (humanized antibodies, single-domain antibody and etc.) were proven to be highly effective for various therapeutic and diagnostic applications [45]. One of the techniques for the production of such antibodies is based on the cloning of the previously identified variable domain sequences from mAbs into the artificially designed vectors encoding constant regions of the antibodies. Therefore, we aimed to determine the amino acid sequence of mAbs #11/9 which it responsible for the binding to S protein. For this reason, we amplified cDNA encoding heavy and light chains of mAbs by 5’ SMART RACE method. Using primers specific to k or λ chains we demonstrated that mAbs #11/9 contains only k light chain (**Fig. 5A**). Sequencing of PCR products revealed the nucleotide and amino acid sequence of variable domains of mAbs #11/9 (**Fig. 5B**). Bioinformatic analysis showed that both light and heavy chains do not contain in frame stop codons, have the correct position of the conserved amino acids and that the corresponding nucleotide sequences have not been published previously. Therefore, the obtained sequences most likely represent viable antibodies originating from murine splenocytes but not abortive rearrangement products which theoretically could appear from X63 myeloma cells that were used for fusion.

**Fig. 5.**
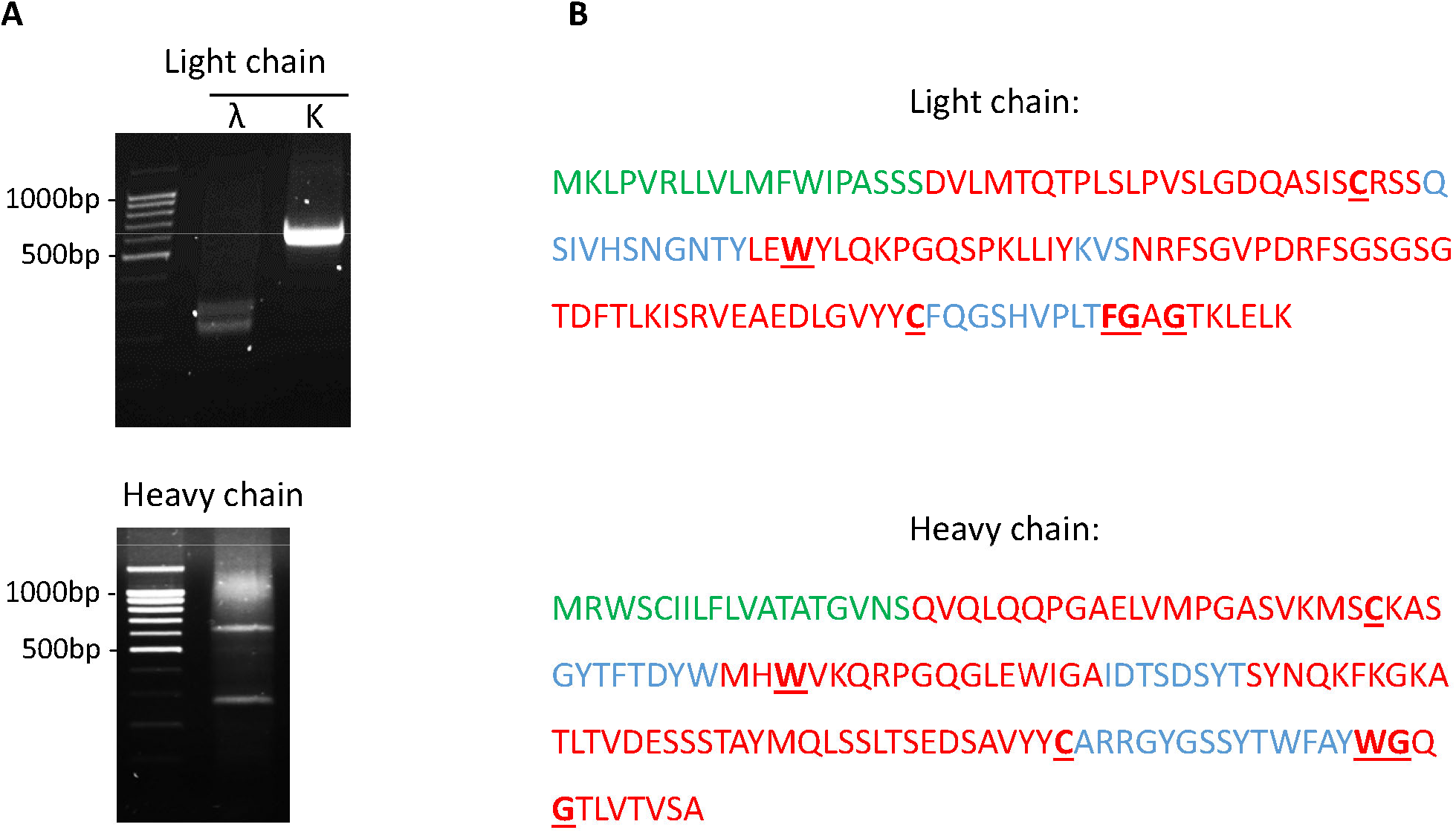
Sequencing of anti-RBD SARS-CoV-2 mAbs. *A*, PCR amplification of cDNA encoding immunoglobulins’ k-chain, λ-chain, and heavy-chain from monoclone #11/9. *B*, Amino acid sequence of variable domains from light and heavy chains of 11/9 antibodies’. Different regions of immunoglobulins are highlighted: Ig leader sequence (green); framework regions (red); complementarity determining regions (blue); conserved amino acids (bold, underlined).

## Discussion

In this study we describe the development of mAbs against RBD fragment of SARS-CoV2 S protein. The expression and purification of the antigen is often the most challenging step of the mAbs production technique. In case of proteins expressed in E. coli the optimization of refolding conditions may frequently represents a major problem and time consuming step. On the other hand, if protein is expressed in mammalian cells it might be difficult to obtain a sufficient amount of the material for animal injection. Here we utilized immunization of mice with insoluble protein mixed with FIA and subsequent reimmunization with cells overexpressing target protein. The main advantage of this protocol is the absence of steps that may require extensive optimization. The only necessary materials are the recombinant protein fragment expressed in E. coli, and the plasmid encoding full length protein. Both components can be easily obtained using standard well established methods. Therefore, we believe that the protocol applied in this study might be useful for the rapid development of mAbs against viral proteins. It became clear during the pandemic of COVID19 that such necessity may indeed arise in the future.

Although the aim of this study was to create mAbs to S protein, we have also obtained a set of data that might be useful for the development of the vaccines against COVID19. Thus, we demonstrated that two injections of RBD mixed with FIA might be sufficient for the emerging of antibodies that can target full length native S protein. On the other hand, it might be worth to note that most of the mAbs that were established during this study could also bind to endogenous protein from human cells. Therefore, further studies are needed to investigate possible cross reactivity of antibodies that arise against SARS-CoV2 S protein.

Another interesting observation could be made during comparison of WBs on **figures 3B** and **4B**. Latter one has much stronger band that corresponds to S protein. Same cells were used for both WBs, however for **figures 4B** we performed cell lysis by boiling in SDS solution, while for WB on **figures 3B** cells were lysed on ice in RIPA buffer. Therefore, the discrepancy in WBs might be attributed to the fact that S protein is tightly anchored to the cell membrane and therefore harsh conditions are required for its’ efficient extraction, while a standard lysis protocol leaves most of S protein in the pellet that contains insoluble cell fragments.

Finally, it is important to note, that for rigorous characterization of the mAbs obtained in this study further experiments are needed. Thus, here we only used culture medium from hybridoma cells or ascitic fluid in which the exact concentration of mAbs was not determined. If the mAbs developed here prove to be useful, it will be important to repeat some of the experiments with purified antibodies taken at various concentrations. This will allow to determine the optimal concentration of mAbs for each application and to calculate the Kd of antibody-antigen binding. Also it will be interesting to repeat experiment with inhibition of cell infection using different experimental models and cell lines. However, we believe that the most useful result of the current study is, firstly, the hybridoma cells that are able to produce mAbs against SARS-CoV2 in any quantity needed and, secondly, the amino acid sequence of mAbs against RBD. Knowledge of these sequence might be useful for the development of the cocktails of neutralizing antibodies against COVID19.

In summary, we developed novel mAbs that might serve as an important tool for the scientific research of SARS-CoV2 and also could help in the establishing of diagnostic and treatment methods for COVID-19 patients. Significant advantage of these mAbs is the presence of corresponding hybridoma cells which could produce these antibodies in nearly any quantity with comparatively little costs.

## Acknowledgments

We thank Dr. Nevan J. Krogan for providing pTwist-EF1a-nCoV-2019-S-2xStrep plasmid. We thank the Center for Precision Genome Editing and Genetic Technologies for Biomedicine, Federal Research and Clinical Center of Physical-Chemical Medicine of Federal Medical Biological Agency for providing RBD fragment of S protein purified from HEK293 cells. This work was supported by The Russian Foundation for Basic Research grants 17-29-06056 (MSP), 18-29-01027 (MIS), 19-34-90102 (TDL), 20-04-00804 (MSP) and Russian Science Foundation grant 19-44-02027 (MSP).

## References

[1] US Centers for Disease Control and Prevention. Coronavirus disease2019(Covid-19): situation summary. https://www.cdc.gov/coronavirus/2019-nCoV/summary.html.

[2] B. Gates, Responding to Covid-19 - A Once-in-a-Century Pandemic? N. Engl. J. Med. 382 (2020) 1677–1679. https://doi.org/10.1056/NEJMp2003762.

[3] Y. Jee, WHO International Health Regulations Emergency Committee for the COVID-19 outbreak. Epidemiol Health. (2020), 42:e2020013. https://doi.org/10.4178/epih.e2020013.

[4] E. Dong, H. Du., L. Gardner, An interactive web-based dashboard to track COVID-19 in real time. Lancet Infect. Dis. 20 (2020) 533–534. https://doi.org/10.1016/S1473-3099(20)30120-1.

[5] R. Lu, X. Zhao, J. Li, P. Niu., B. Yang, Genomic characterisation and epidemiology of 2019 novel coronavirus: Implications for virus origins and receptor binding. Lancet. 6736 (2020) 1–10. https://doi.org/10.1016/S0140-6736(20)30251-8.

[6] A. Roy, A. Kucukural, Y. Zhang, I-TASSER: a unified platform for automated protein structure and function prediction. Nature Protocols. 5 (2010) 725–738. https://doi.org/10.1038/nprot.2010.5.

[7] A. E. Gorbalenya, S. C. Baker, R. S. Baric, R. J. de Groot, C. Drosten, A. A. Gulyaeva, B. L. Haagmans, C. Lauber, A. M. Leontovich, B. W. Neuman, D. Penzar, S. Perlman, L. L. M. Poon, D. V. Samborskiy, I. A. Sidorov, I. Sola, J. Ziebuhr, The species Severe acute respiratory syndrome-related coronavirus: classifying 2019-nCoV and naming it SARS-CoV-2. Nat Microbiol. 5 (2020) 536–544. https://doi.org/10.1038/s41564-020-0695-z.

[8] T. Phan, Genetic diversity and evolution of SARS-CoV-2. Infect Genet Evol. 81 (2020), 104260. https://doi.org/10.1016/j.meegid.2020.104260.

[9] D. Mercatelli, F. M. Giorgi, Geographic and Genomic Distribution of SARS-CoV-2 Mutations. Front. Microbiol. 11 (2020) 1800. https://doi.org/10.3389/fmicb.2020.01800.

[10] F. Li, W. Li, M. Farzan, S. C. Harrison, Structure of SARS coronavirus spike receptor-binding domain complexed with receptor. Science. 309(2005)1864–1868. https://doi.org/10.1126/science.1116480.

[11] P. Zhou, X.-L. Yang, X.-G. Wang, B. Hu, L. Zhang, W. Zhang, H.-R. Si, Y. Zhu, B. Li, C.-L. Huang, H.-D. Chen, J. Chen, Y. Luo, H. Guo, R.-D. Jiang, M.-Q. Liu, Y. Chen, X.-R. Shen, X. Wang, X.-S. Zheng, K. Zhao, Q.-J. Chen, F. Deng, L.-L. Liu, B. Yan, F.-X. Zhan, Y.-Y. Wang, G. Xiao, Z.-L. Shi, Discovery of a novel coronavirus associated with the recent pneumonia outbreak in humans and its potential bat origin. BioRxiv. 579 (2020) 270–273. https://doi.org/10.1101/2020.01.22.914952.

[12] M. Letko, A. Marzi, V. Munster, Functional assessment of cell entry and receptor usage for SARS-CoV-2 and other lineage B betacoronaviruses. Nat. Microbiol. 5 (2020) 562–569. https://doi.org/10.1038/s41564-020-0688-y.

[13] G. Lu, Y. Hu, Q. Wang, J. Qi, F. Gao, Y. Li, Y. Zhang, W. Zhang, Y. Yuan, J. Bao, B. Zhang, Y. Shi, J. Yan, G. F. Gao, Molecular basis of binding between novel human coronavirus MERS-CoV and its receptor CD26. Nature. 500 (2013) 227–231. https://doi.org/10.1038/nature12328.

[14] D. Wrapp, N. Wang, K. S. Corbett, J. A. Goldsmith, C. Hsieh, O. Abiona, B. S. Graham, J. S. McLellan, Cryo-EM structure of the 2019-nCoV spike in the prefusion conformation. Science 367 (2020) 1260–1263. https://doi.org/10.1126/science.abb2507.

[15] J. Lan, J. Ge, J. Yu, S. Shan, H. Zhou, S. Fan, Q. Zhang, X. Shi, Q. Wang, L. Zhang, X. Wang, Structure of the SARS-CoV-2 spike receptor-binding domain bound to the ACE2 receptor. Nature 581 (2020) 215–220. https://doi.org/10.1038/s41586-020-2180-5.

[16] J. Shang, G. Ye, K. Shi, Y. Wan, C. Luo, H. Aihara, Q. Geng, A. Auerbach, F. Li, Structural basis of receptor recognition by SARS-CoV-2. Nature. 581 (2020) 221–224. https://doi.org/10.1038/s41586-020-2179-y.

[17] A. M. Baig, A. Khaleeq, S. Hira, Elucidation of cellular targets and exploitation of the receptor binding domain of SARS-CoV-2 for vaccine and monoclonal antibody synthesis. Journal of Medical Virology. (2020) 1–12. https://doi.org/10.1002/jmv.26212.

[18] R. Channappanavar, C. Fett, J. Zhao, D. K. Meyerholz, S. Perlman, Virus-specific memory CD8 T cells provide substantial protection from lethal severe acute respiratory syndrome coronavirus infection. J. Virol. 88 (2014) 11034–11044. https://doi.org/10.1128/JVI.01505-14.

[19] L. Ni, F. Ye, M.-L. Cheng, Y. Feng, Y.-Q. Deng, H. Zhao, P. Wei, JiwanGe, MengtingGou, X. Li, L. Sun, T. Cao, P. Wang, C. Zhou, R. Zhang, P. Liang, H. Guo, X. Wang, C. Dong, Detection of SARS-CoV-2-specific humoral and cellular immunity in COVID-19 convalescent individuals. Immunity. 52 (2020) 971–977.e3. https://doi.org/10.1016/j.immuni.2020.04.023.

[20] N. Lurie, M. Saville, R. Hatchett, J. Halton, Developing Covid-19 vaccines at pandemic speed. N Engl J Med. 382 (2020) 1969–1973. https://doi.org/10.1056/NEJMp2005630.

[21] Coronavirus Vaccine Tracker https://www.nytimes.com/interactive/2020/science/coronavirus-vaccine-tracker.html (accessed 21 August 2020).

[22] H. Shi, X. Han, N. Jiang, Y. Cao, O. Alwalid, J. Gu, Y. Fan, C. Zheng, Radiological findings from 81 patients with COVID-19 pneumonia in Wuhan, China: a descriptive study. Lancet Infect Dis. 20 (2020) 425–434. https://doi.org/10.1016/S1473-3099(20)30086-4.

[23] A. Bernheim, X. Mei, M. Huang, Y. Yang, Z. A. Fayad, N. Zhang, K. Diao, B. Lin, X. Zhu, K. Li, S. Li, H. Shan, A. Jacobi, M. Chung, Chest CT findings in coronavirus disease-19 (COVID-19): relationship to duration of infection. Radiology. 295(2020), 200463. https://doi.org/10.1148/radiol.2020200463.

[24] L. Guo, L. Ren, S. Yang, M. Xia, D. Chang, F. Yang, C. S Dela Cruz, Y. Wang, C. Wu, Y. Xiao, L. Zhang, L. Han, S. Dang, Y. Xu, Q.-W. Yang, S.-Y. Xu, H.-D. Zhu, Y.-C. Xu, Q. Jin, L. Sharma, L. Wang, J. Wang, Profiling early humoral response to diagnose novel coronavirus disease (COVID-19). Clin Infect Dis. 71 (2020) 778–785. https://doi.org/10.1093/cid/ciaa310.

[25] J. Zhao, Q. Yuan, H. Wang, W. Liu, X. Liao, Y. Su, X. Wang, J. Yuan, T. Li, J. Li, S. Qian, C. Hong, F. Wang, Y. Liu, Z. Wang, Q. He, Z. Li, B. He, T. Zhang, Y. Fu, S. Ge, L. Liu, J. Zhang, N. Xia, Z. Zhang, Antibody responses to SARS-CoV-2 in patients of novel coronavirus disease 2019. Clin Infect Dis. (2020), ciaa344. https://doi.org/10.1093/cid/ciaa344.

[26] T. F. Rogers, F. Zhao, D. Huang, N. Beutler, A. Burns, W.-T. He, O. Limbo, C. Smith, G. Song, J. Woehl, L. Yang, R. K. Abbott, S. Callaghan, E. Garcia, J. Hurtado, M. Parren, L. Peng, S. Ramirez, J. Ricketts, M. J. Ricciardi, S. A. Rawlings, N. C. Wu, M. Yuan, D. M. Smith, D. Nemazee, J. R. Teijaro, J. E. Voss, I. A. Wilson, R. Andrabi, B. Briney, E. Landais, D. Sok, J. G. Jardine, D. R. Burton, Isolation of potent SARS-CoV-2 neutralizing antibodies and protection from disease in a small animal model. Science. 369 (2020) 956–963. https://doi.org/10.1126/science.abc7520.

[27] W. Tai, X. Zhang, Y. He, S. Jiang, L. Du, Identification of SARS-CoV RBD-targeting monoclonal antibodies with cross-reactive or neutralizing activity against SARS-CoV-2. Antiviral Res. 179 (2020), 104820. https://doi.org/10.1016/j.antiviral.2020.104820.

[28] A. Baum, B. O. Fulton, E. Wloga, R. Copi, K. E. Pascal, V. Russo, S. Giordano, K. Lanza, N. Negron, M. Ni, Y. Wei, G. S. Atwal, A. J. Murphy, N. Stahl, G. D. Yancopoulos, C. A. Kyratsous, Antibody cocktail to SARS-CoV-2 spike protein prevents rapid mutational escape seen with individual antibodies. Science. 369 (2020) 1014–1018. https://doi.org/10.1126/science.abd0831.

[29] H. Kaplon, M. Muralidharan, Z. Schneider, J. M. Reichert, Antibodies to watch in 2020. MAbs. 12 (2020), 1703531. https://doi.org/10.1080/19420862.2019.1703531.

[30] A. O. Hassan, J. B. Case, E. S. Winkler, L. B. Thackray, N. M. Kafai, A. L. Bailey, B. T. McCune, J. M. Fox, R. E. Chen, W. B. Alsoussi, J. S. Turner, A. J. Schmitz, T. Lei, S. Shrihari, S. P. Keeler, D. H. Fremont, S. Greco, P. B. McCray, S. Perlman, M. J. Holtzman, A. H. Ellebedy, M. S. Diamond, A SARS-CoV-2 Infection Model in Mice Demonstrates Protection by Neutralizing Antibodies. Cell. 182 (2020) 744–753.e4. https://doi.org/10.1016/j.cell.2020.06.011.

[31] D. E. Gordon, G. M. Jang, M. Bouhaddou, J. Xu, K. Obernier, K. M. White, M. J. O’Meara, V. V. Rezelj, J. Z. Guo, D. L. Swaney, A SARS-CoV-2 protein interaction map reveals targets for drug repurposing. Nature. 583 (2020) 459–468. https://doi.org/10.1038/s41586-020-2286-9.

[32] H.-Y. Kim, A. Stojadinovic, M. J. Izadjoo, Immunization, Hybridoma Generation, and Selection for Monoclonal Antibody Production. Methods Mol Biol. 1131 (2014) 33–45. https://doi.org/10.1007/978-1-62703-992-5_3.

[33] N. Rose, C. A. Pinho-Nascimento, A. Ruggieri, P. Favuzza, M. Tamborrini, H. Roth, M. T. B. de Moraes, H. Matile, T. Jänisch, G. Pluschke, K. Röltgen, Generation of monoclonal antibodies against native viral proteins using antigenexpressing mammalian cells for mouse immunization. BMC Biotechnol. 16 (2016), 83. https://doi.org/10.1186/s12896-016-0314-5.

[34] M. S. Pavlyukov, N. V. Antipova, M. V. Balashova, T. V. Vinogradova, E. P. Kopantzev, M. I. Shakhparonov, Survivin monomer plays an essential role in apoptosis regulation. J Biol Chem. 286 (2011) 23296–23307. https://doi.org/10.1074/jbc.M111.237586.

[35] A. L. Tarentino, T. H. Plummer, Enzymatic deglycosylation of asparagine-linked glycans: Purification, properties, and specificity of oligosaccharide-cleaving enzymes from Flavobacterium meningosepticum. Guide to Techniques in Glycobiology. 230 (1994) 44–57. https://doi.org/10.1016/0076-6879(94)30006-2.

[36] M. S. Pavlyukov, H. Yu, S. Bastola, M. Minata, V. O. Shender, Y. Lee, S. Zhang, J. Wang, S. Komarova, J. Wang, S. Yamaguchi, H. A. Alsheikh, J. Shi, Do. Chen, A. Mohyeldin, S.-H. Kim, Y. J. Shin, K. Anufrieva, E. G. Evtushenko, N. V. Antipova, G. P. Arapidi, V. Govorun, N. B. Pestov, M. I. Shakhparonov, L. J. Lee, D.-H. Nam, I. Nakano, Apoptotic Cell-Derived Extracellular Vesicles Promote Malignancy of Glioblastoma Via Intercellular Transfer of Splicing Factors. Cancer Cell. 34 (2018) 119–135. https://doi.org/10.1016/j.ccell.2018.05.012.

[37] X. Wang, M. McManus, Lentivirus Production. J. Vis. Exp. 32 (2009), e1499. https://doi.org/10.3791/1499.

[38] L. Meyer, T. López, R. Espinosa, C. F. Arias, C. Vollmers, R. M. DuBois, A simplified workflow for monoclonal antibody sequencing. PloS one. 14 (2019), e0218717. https://doi.org/10.1371/journal.pone.0218717

[39] C.-C. Huang, K.-W. Cheng, Y.-C. Hsieh, W.-W. Lin, C.-M. Cheng, S.-S. F. Yuan, I-J. Chen, Y.-A. Cheng, Y.-C. Lu, B.-C. Huang, Y.-C. Tung, T.-L. Cheng, Use of syngeneic cells expressing membrane-bound GM-CSF as an adjuvant to induce antibodies against native multi-pass transmembrane protein. Sci Rep. 9 (2019), 9931. https://doi.org/10.1038/s41598-019-45160-9.

[40] S. Wang, S. Lu, DNA immunization. Current protocols in microbiology. 31 (2013) 18.3.1–18.3.24. https://doi.org/10.1002/9780471729259.mc1803s31.

[41] W. Viranaicken, B. Nativel, P. Krejbich-Trotot, W. Harrabi, S. Bos, C. E. Kalamouni, M. Roche, G. Gadea, P. Despres, ClearColi BL21(DE3)-based expression of Zika virus antigens illustrates a rapid method of antibody production against emerging pathogens. Biochimie. 142 (2017) 179–182. https://doi.org/10.1016/j.biochi.2017.09.011.

[42] F. J. Castillo, L. J. Mullen, B. C. Grant, J. DeLeon, J. C. Thrift, L. W. Chang, J. M. Irving, D. J. Burke, Hybridoma stability. Dev Biol Stand. 83 (1994) 55–64.

[43] N. Vankadari, J. A. Wilce, Emerging WuHan (COVID-19) coronavirus: glycan shield and structure prediction of spike glycoprotein and its interaction with human CD26. Emerg Microbes Infect. 9 (2020) 601–604. https://doi.org/10.1080/22221751.2020.1739565.

[44] G. Simmons, J. D. Reeves, A. J. Rennekamp, S. M. Amberg, A. J. Piefer, P. Bates, Characterization of severe acute respiratory syndrome-associated coronavirus (SARS-CoV) spike glycoprotein-mediated viral entry. Proc Natl Acad Sci USA. 101 (2004) 4240–4245. https://doi.org/10.1073/pnas.0306446101.

[45] N. C. Peterson, Advances in Monoclonal Antibody Technology: Genetic Engineering of Mice, Cells, and Immunoglobulins. ILAR Journal. 46 (2005) 314–319. https://doi.org/10.1093/ilar.46.3.314.

